# Evaluation of saliva as a source of accurate whole-genome and microbiome sequencing data

**DOI:** 10.1101/2020.11.16.384438

**Authors:** Anthony F. Herzig, Lourdes Velo-Suárez, Gaёlle Le Folgoc, Anne Boland, Hélène Blanché, Robert Olaso, Liana Le Roux, Christelle Delmas, Marcel Goldberg, Marie Zins, Franck Lethimonnier, Jean-François Deleuze, Emmanuelle Génin

## Abstract

This study sets out to establish the suitability of saliva-based whole-genome sequencing (WGS) through a comparison against blood-based WGS. To fully appraise the observed differences, we developed a novel technique of pseudo-replicates. We also investigated the potential of characterising individual salivary microbiomes from non-human DNA fragments found in saliva.

We observed the majority of discordant genotype calls between blood and saliva calls fell into known regions of the human genome that are typically sequenced with low confidence; and could be identified by quality control measures. Pseudo-replication demonstrated that the levels of discordance between blood- and saliva-derived WGS data were entirely similar to what one would expect between technical replicates if an individual’s blood or saliva had been sequenced twice. Finally, we successfully sequenced salivary microbiomes in parallel to human genomes as demonstrated by a comparison against the Human Microbiome Project.

**Author Summary:** DNA is usually collected from blood for the analysis of human genomes. In France, a new and very large genetic dataset will be created where selected participants will be sent saliva-collection kits in the post as this data collection method presents numerous logistical benefits. It has been previously shown that good quality genetic data can be created from saliva, though existing studies have often not considered the latest technologies or have only analysed a very small number of individuals. In this study, we have analysed genetic data derived from saliva for 39 individuals to give a firm conclusion that the proposed genome sequencing approach of the new French dataset will be capable of provided high quality data by making a comparison to pre-existing genetic data derived from blood for these 39 individuals. In order to do so, we developed a novel method (presented here) to establish the similarity between two sets of genetic data for the same individual that are generated from separate DNA samples. Finally, we have also demonstrated an added bonus of colleting saliva samples: that it is possible to gather both human genetic data and potentially interesting salivary microbiome data at the same time by separating and analysing in parallel human and non-human DNA fragments.

## Introduction

Whole-genome sequencing in humans has become widespread in the exploration of genetic variation between and within populations as well as in genetic epidemiology. Reduced costs have made such sequencing far more attainable. For the majority of large panels of human genetic variation, for example those that have been gathered to for the Haplotype Reference Consortium [1], genomic data has been generated through whole-genome sequencing (WGS) of DNA extracted from blood. Quoted error rates for most WGS technologies are generally very low [2], yet even apparently tiny error rates can represent hundreds and thousands of erroneous calls across entire genomes. Errors may well congregate in regions that are particularly complicated to sequence as the accuracy of WGS has been shown to vary in different regions of the genome [3–5]. Hence, when planning the creation of a new large whole-genome sequencing dataset using a methodology that is in some sense alternative, it is vital to establish that it does not introduce unexpected and problematic patterns into the data that would render the new dataset incomparable to existing WGS datasets.

Saliva has many advantages over blood in terms of the logistics of data collection and individual participation levels [6,7]. The POPGEN project in France envisages the creation of a French genomic reference panel by gathering DNA samples using Oragene OG-600 saliva kits. These are to be sent out, completed by participants and returned through the post. A similar approach was used in a recent study in the USA which took advantage of both social media and the use of saliva kits to increase the engagement with vast numbers of individuals [8], and to facilitate the creation of a large population based study with individuals spread evenly across a large region. This last point is of great importance if a full representation of a population’s genetics is to be recorded and also to avoid potential geographical selection biases that have recently been highlighted by Haworth et al. [9] and by Munafò et al. [10] in the ALSPAC cohort [11] and in the UK Biobank [12], respectively.

DNA extraction from saliva is by no means a new concept, and has been demonstrated to be a successful technique for genotyping and sequencing studies. The quality and quantity of attained DNA had been shown to be superior in saliva as compared to buccal swabs [6,13,14], likely due to the higher prevalence of leukocytes in saliva [15]. Limiting factors for saliva for sequencing were however presented in [16] where the presence of non-human DNA and subsequent low quantities of attained human DNA was postulated as resulting in large observed differences in the number of markers that could be called (significantly less in saliva than in blood); this being an observation that also made by Herráez and Stoneking [17]. Indeed, even in a recent study that compared WGS data between blood and saliva [18], the most problematic characteristics regarding data from saliva included a large number of samples failing an initial agarose gel quality control test (suggesting very poor quality DNA) and the potential for the presence of a large proportion of non-human DNA. The possibility for low quality [19] and low quantity [20] of human DNA in certain saliva samples has also been presented in the sequencing data generated via exome-capture kits.

At time of writing, comparisons of genomic data between samples of DNA from blood and saliva have largely concluded that saliva is an adequate substitute for blood; producing data with high but not perfect concordance to genotype calls from blood [6,21–28]. If concordance rates of the order of 95-99% between genotypes obtained from blood and saliva samples are to be considered satisfactory, it is important to realize that at the scale of a whole genome sequence where ~3 million genetic variants are expected per individual, this could represent as many as 150,000 differences. Such reported statistics, we felt, are worrisome considering they could predict very high error rates in saliva-based sequencing data and could predict unwanted artefacts that could significantly inhibit the utility of a large population-wide WGS dataset. Furthermore, many of the current publications that compare blood against saliva have employed very small sample sizes (<10) and few have examined WGS data.

In this study, we compare WGS data derived from blood and saliva DNA from 39 individuals. Without gold-standard genomes to evaluate the quality of the sequencing in GAZEL-ADN, we have leveraged information on intrinsic difficulties in sequencing certain regions of the genome set out in a collection of studies from the Genome in a Bottle (GiaB) project [3,29,30]. Furthermore, we introduce a novel technique of pseudo-replication. This is a data-driven method to generate, in the absence of gold standard genomes, specific baselines for sequencing reproducibility from within a study. We also show that whilst the treatment of our samples followed the protocol of sequencing human DNA rather than microbiome DNA, we were able to sequence and identify large numbers of non-human DNA fragments in order to give a characterisation of individual salivary-microbiomes in our sample of 39 individuals. This presents an added bonus of the experimental design of using saliva-kits for data collection as one can sequence both good quality human and microbiome data in parallel.

## Results

### Comparison of blood and saliva derived WGS data at the individual level

Details of the recruitment of individuals, the DNA extraction, sequencing, alignment, and genotype calling for blood and saliva are given in the Methods. We examined similarities between single sample WGS variant calls from blood and saliva for each of the 39 individuals. In Supplementary Figure 1, the number of called variants, the mean individual read depths, and the mean genotype quality scores are presented. We also calculated per-individual statistics from BAM files; Supplementary Figure 1 includes results for mean insert length of reads, estimated base error rates, and mean read quality. We saw similar numbers of variants being called (mean of 4,619,812 for saliva against a mean of 4,615,935 for blood) but higher statistics regarding the depth and quality in saliva. In saliva the mean read depth was 40.2× (range of 28.2× to 54.2×) compared to 35.0× for blood (range of 29.2× to 39.0×). Following a similar trend, the mean Genotype Quality (GQ) observed in saliva was 95.3 (range of 90.2 to 97.0) compared to 93.5 (range of 89.8 to 95.2) for blood. Within individual BAM files, we observed that saliva samples had longer average insert lengths, lower estimated error rates, and higher average read quality scores. Between each pair of variant call files (for each individual) the mean percentage of sites that overlap (matched for chromosome, position and both reference and alternative alleles) out of the total number of variants observed in either blood or saliva was 96.3% (range 95.8% to 96.7%). On average per-individual, 86,591 variants are only found in the WGS data coming from saliva against 82,714 found only in WGS data from blood. For the variants in common between the two call sets, on average 99.6% of the calls agree; with higher agreement for SNPs (mean of 99.8%) compared to INDELs (98.4%) (Supplementary Figure 2). These summary statistics indicate that the quality of the data derived from saliva is here above that of blood. This is not a generalizable observation, simply a characteristic of our particular study likely related to the fact that different genome sequencers were used for blood and saliva (see Methods).

### Comparison of blood and saliva derived WGS data at the whole study level

Joint sample calling was performed for blood and saliva for all 39 samples, giving a variant call file (VCF) with 78 columns of WGS data that contains 12,085,848 bi-allelic variants across the 22 autosomal chromosomes. The differences in observable quality in the single sample calls translated here to small differences in levels of genotype missingness (per individual) between saliva and blood (mean of 0.006 for saliva against 0.011 for blood). For each polymorphic genetic variant present in the joint-called VCF, we calculated the F-measure [31,32] (also known as the Sørensen-Dice coefficient, see Methods) between the 39 saliva- and blood-based genotypes. The F-measure ranges between 0 and 1, with a value of 1 indicating complete agreement between calls from blood and saliva and lower values indicating diminished agreement of variant calls across the 39 individuals. Discordances between blood and saliva genotypes can be assumed to indicate genotyping errors in at least one of the two datasets as we should not anticipate any significant contribution from somatic mosaicism between saliva and blood at this depth of sequencing [33].

In Figure 1(a), the distribution of the F-measures is displayed for all 12,085,848 variants. 93.61% of all variants show perfect agreement between blood and saliva. Subsequently in Figure 1(a), we split these variants into two genomic region-sets: high-confidence regions indicated by the GiaB project and the complementing set of low-confidence regions. Subsequently, we focused only on the variants that pass our quality control (QC) thresholds (see Methods). The GiaB high-confidence region list was accessed from ftp://ftp-trace.ncbi.nlm.nih.gov/giab/ftp/release. We observed that a high proportion of the variants with disagreements between blood and saliva fall within the low-confidence regions of the genome. It is also clear that quality control (described fully in the Methods) was successful at removing discordant variants in both high and low-confidence regions. In Figure 1(b), we plot the F-measure along chromosome 2 as an example of how discordant variants cluster into the low-confidence GiaB regions and how quality control improves concordance. Similar patterns are observed over all chromosomes (Supplementary Figure 3).

**Figure 1.**
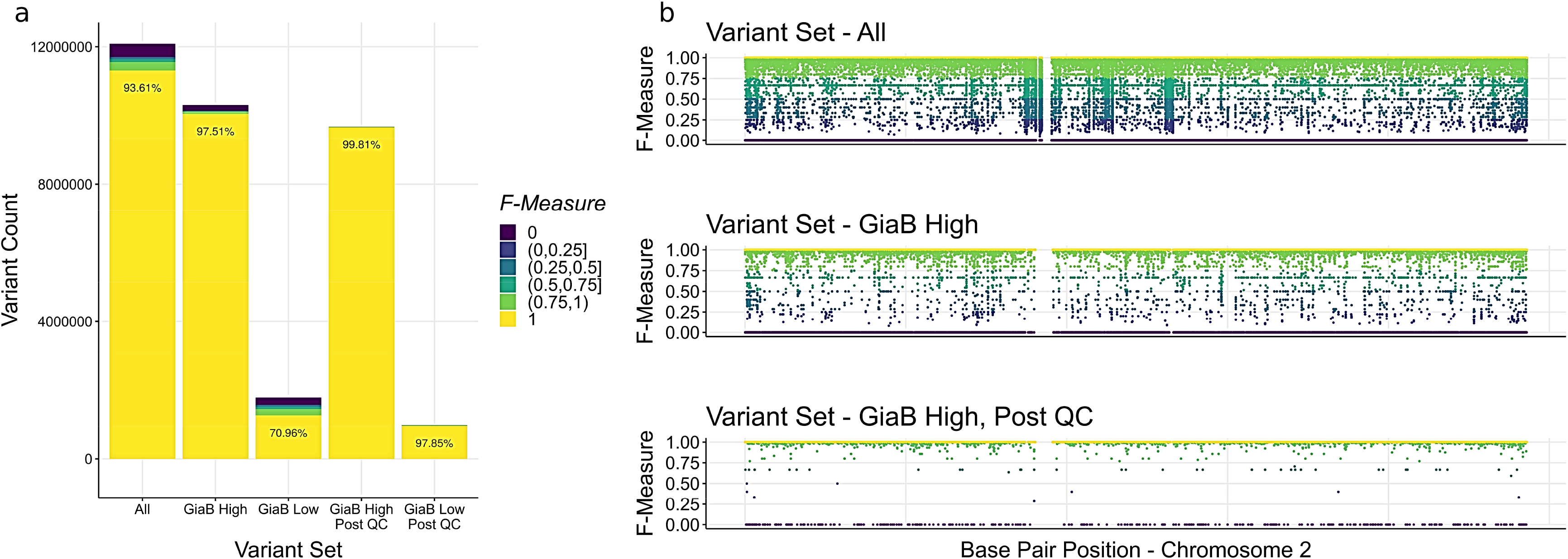
Per-variant F-measure statistics between blood and saliva under different filtering conditions. (a) For five different variant sets, the distribution of the F-measure (harmonic mean of precision and recall) is given. Yellow represents the variants that are 100% concordant (F-measure of 1) between the 39 blood and 39 saliva WGS call sets. The percentages of variants with an F-measure of 1 are overlaid for each variant set. Darker colours represent lower intervals of the F-measure. The first variant set is all variants observed across the 22 autosomal chromosomes (All 12,085,848 variants). The second and third set are complementary, representing the high- and low-confidence regions of the genome as indicated by the Genome in a Bottle (GiaB) project (10,308,126 ‘GiaB High’ variants and 1,777,722 ‘GiaB Low’ variants). The fourth and fifth sets show distribution of the F-measure again in high and low-confidence regions but after applying qualitycontrol thresholds (9,679,699 ‘GiaB high – Post QC’ variants and 983,381 ‘GiaB Low – Post QC’ variants). (b) For the variant sets ‘All’, ‘GiaB High’ and ‘GiaB High – Post QC’, the F-measures from chromosome 2 are plotted against base-pair positions to give an illustration of the congregation of poorly concordant variants in low-confidence regions and also the efficacy of quality control.

### Pseudo-replication

We have ascertained that the majority of the discordances observed between saliva and blood could be segregated using quality control and the GiaB low-confidence region list. This gave a strong suggestion that the differences between blood and saliva that we observed were in the majority due to readily identifiable sequencing errors. This suggested the saliva-based sequencing produced high quality genotype calls (as demonstrated by the high concordance with blood-based sequencing in the high-confidence regions). The success in removing a large proportion of the erroneous calls using quality control also demonstrated that it should not be problematic to analyse saliva- and blood-based WGS data side-by-side.

Yet even after quality control, discordances remain between blood and saliva across our 39 individuals. The question remains as to whether the remaining discordance was linked to the differences between saliva and blood and the respective sequencing pipelines, or falls within the range that would be expected were an individual’s DNA to be sequenced twice on the same platform.

To answer this question, ideally, we would re-sequence each individual’s blood DNA sample in order to compare discrepancies of blood against blood with blood against saliva. However, such resequencing was not possible in this study and could well have led to further batch effects given that there would have been a significant length of time between sequencing runs. Not having access to repeated sequencing data, we decided on an innovative in silico approximation of such a round of re-sequencing and to create specific baselines for the comparisons of variant calling between saliva and blood. This pseudo-replication process is described fully in the Methods. The approach involved returning to the raw FASTQ files which contain lists of each individual’s raw read data from the sequencer. These lists of reads for blood and saliva were each divided into two non-overlapping groups to give four separate lists of raw reads for each individual. These four lists were then processed separately and identically to produce four variant call sets for each individual; two from blood and two from saliva.

Having four variant call sets for each individual allowed us to make six pairwise comparisons; two comparisons between pairs of pseudo replicates derived from either both from blood or saliva (‘blood – blood’ and ‘saliva – saliva’, respectively), and four comparisons between blood and saliva. The same quality control criteria were applied to the 156 (39×4) pseudo-replicates as had been applied to the 78 (39×2) WGS samples previously. For a pair of pseudo-replicates, we calculated a single F-measure across the 22 autosomal chromosomes.

In Figure 2 we present boxplots of the F-measures for the three potential configurations of pairwise comparisons. For each individual, we selected one of the four possible ‘blood – saliva’ comparisons possible in order that each boxplot consists of 39 F-measures. Results given split between high- and low-confidence GiaB regions and were calculated post QC. When concentrating on the sets of variants that are in high-confidence regions (Figure 2), we observed near perfect agreement (F-measures close to 1). In both high- and low-confidence regions, there was no clear separation between the different types of pairwise comparison for each variant set. The ‘blood – blood’ and ‘saliva – saliva’ comparisons provide benchmarks for reproducibility of WGS in our study; serving as approximations of the F-measures that would have been seen if the same individuals had been sequenced twice (but at lower depth). Hence as the ‘blood – saliva’ comparisons gave F-measures in the same range, the differences in tissue, sequencing machines, and other steps of the data preparation did not result in a batch effect between our two sets of WGS data. Furthermore, this demonstrated that WGS of saliva leads to very similar accuracy as WGS of blood. Slightly higher F-statistics for the ‘saliva – saliva’ comparisons (blue boxplots in Figure 2) reflect the slightly higher sequencing quality in saliva in this study.

**Figure 2.**
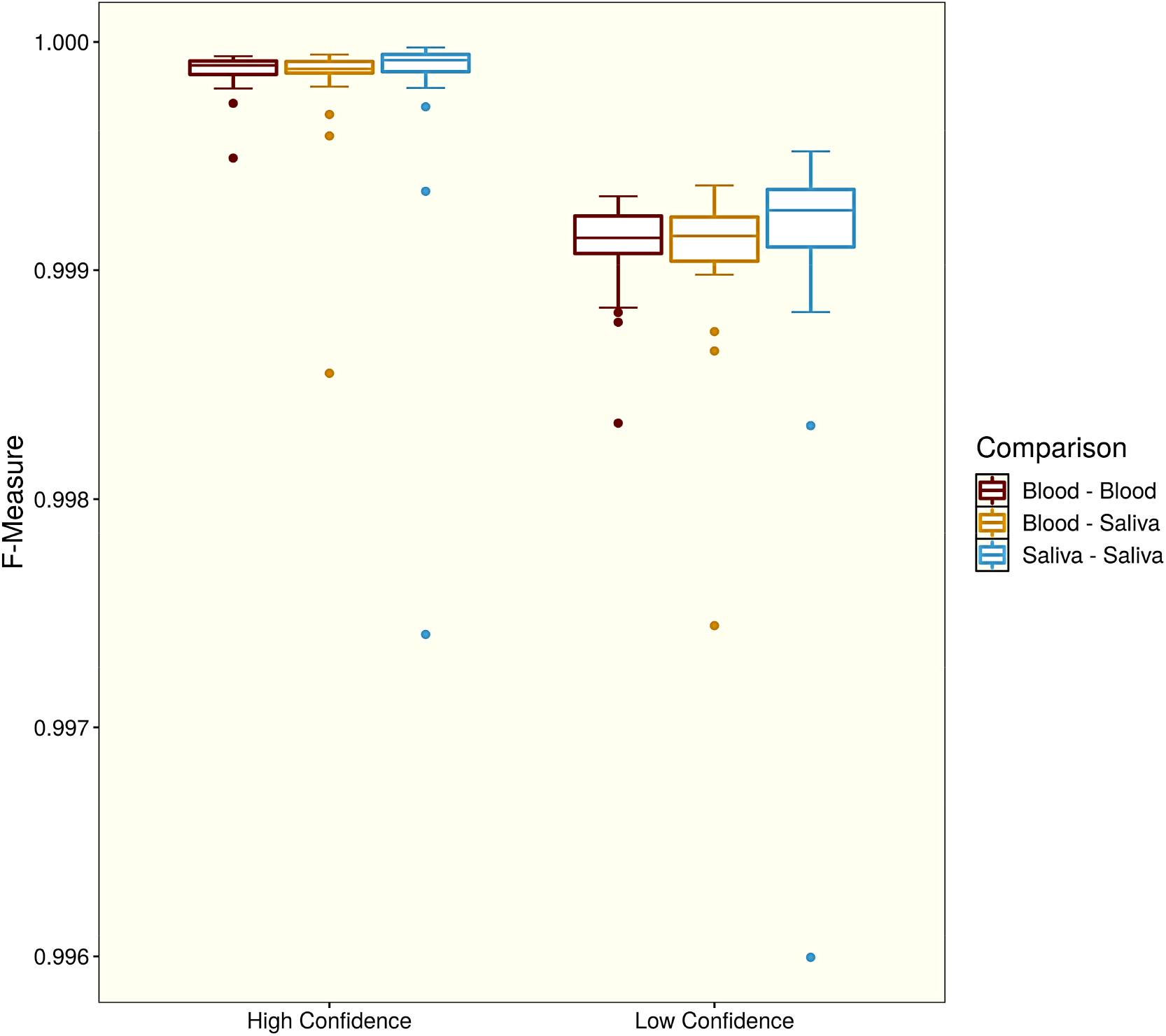
F-measures between pseudo-replicates to mimic a resequencing study. For all 39 individuals with paired blood- and saliva-based WGS data, four variant call sets were generated by creating two pseudo-replicates for both blood and saliva. For each pair of blood- and saliva-based WGS data, three comparisons were made, one between the two blood pseudo-replicates (red), one between a randomly chosen blood pseudo-replicate and a randomly chosen saliva pseudo-replicates (gold), and one between the two saliva pseudo-replicates (blue). The F-measures for these comparisons were calculated for variants in high- and low-confidence regions of the genome after quality control had been applied to all 78 pseudo replicates (left and right, respectively).

A single individual was a distinctive outlier in the analysis of Figure 2 for the ‘saliva – saliva’ comparisons (lowest blue points on 3^rd^ and 6^th^ boxplots). The low F-statistics for this individual suggests a higher number of genotyping errors in the two saliva pseudo-replicates. Though the individual did not have the lowest genomic coverage or average genotype quality among the saliva samples, the quantity of DNA extracted for the individual was close to the threshold for exclusion DNA (only 34.0 ng/μL compared to an average of 63.4 ng/μL across all individuals). Retracing the steps of the DNA extraction period of GAZEL-ADN, we noticed that this individual’s sample was the last to arrive and was processed the same day that their saliva sample arrived in the post. Therefore, the recommendation of Maxwell (PROMEGA) that storing the collectors containing saliva samples for more than two days after collection may improve extraction was not be met and so could explain the slightly lower F-measure involving this individual in Figure 2.

### Salivary Microbiome

We constructed a pipeline to investigate the possibility of sequencing salivary microbiomes from the raw sequencing data derived from saliva. This was necessitated as our raw data had been generated using technology designed for sequencing human DNA while studies of microbiome data typically involve different molecular sequencing approaches [34]. Our goal was to identify reads that were not of human origin and to map such reads to known bacterial reference genomes. The pipeline we used is described fully in the Methods. Applying this to the 39 individuals of the pilot study, we successfully captured an average of 40 million reads that passed quality control measures and could be identified as non-human (Supplementary Figure 4).

These groups of non-human reads were then aligned to known bacterial 16S rRNA gene reference libraries. Across all 39 individuals we qualified a specific taxon to be present in GAZEL-ADN if more than 50 reads aligned to that taxon’s reference 16s ribosomal gene. The OTU groups in GAZEL-ADN were compared to 290 publicly available salivary microbiome samples of the Human Microbiome Project (HMP) [35] (Figure 3a). The majority of salivary phyla that are strongly represented in HMP (nodes close to the center of the taxonomic tree) were found in GAZEL-ADN and only a small proportion of rare genera present in the HMP data were not detectable in GAZEL-ADN reads (grey nodes). These results show that the salivary microbiomes characterized in this study have realistic profiles as no major taxonomy groups from the HMP are missing. Furthermore, many phylogenetic families and genera that were detected in GAZEL-ADN were in fact not present in the HMP data (taxonomic tree edges in blue). This might suggest that the deep shotgun sequencing performed in GAZEL-ADN was potentially more sensitive than the 16S analysis of the HMP. However, it is not possible to compare the relative proportions of reads in each OTU group between GAZEL-ADN and the HMP due to the differences in DNA amplification methods and the fact that meta-genomic microbiome studies cannot be assumed to reflect a linear transformation of true microbial population sizes in the body [36,37]. Hence, these results should be interpreted with caution. The proportions of reads that match to the main phyla presented in Figure 3a (11 branches that lead out from the center) for each individual of GAZEL-ADN are presented in Figure 3b where notable difference between individuals can be observed.

**Figure 3.**
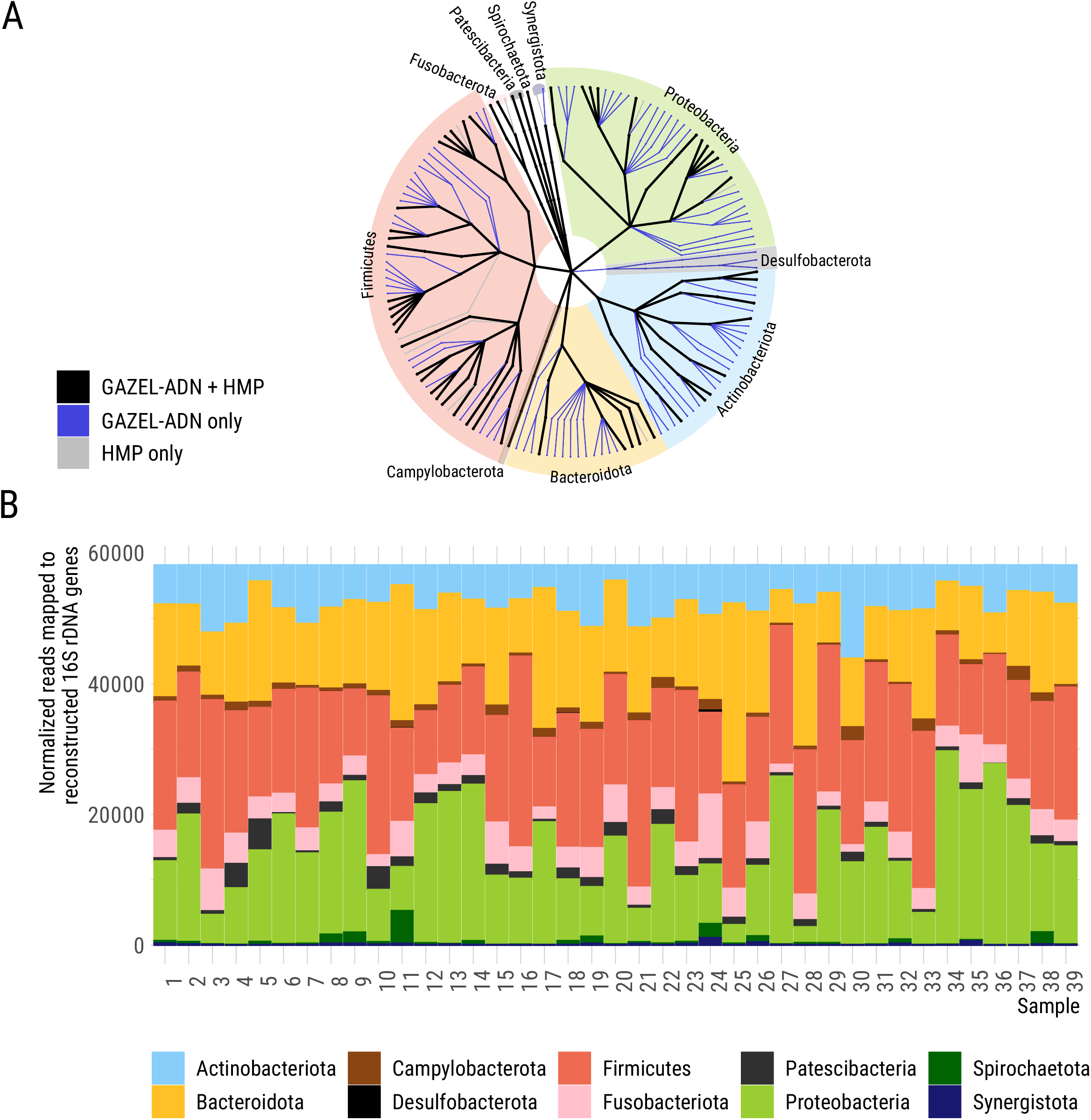
Analysis of the salivary microbiome. (a) Phylogenetic tree of the Operational taxonomic unit (OTU) assignments of non-human reads in GAZEL-ADN. Black nodes represent taxonomic groups (Kingdom, Phylum, Class, Order, Family and Genus from the center to the edge) that were present in both GAZEL-ADN and the publicly available data on salivary microbiomes obtained from 16S NGS analysis from the Human Microbiome Project (HMP). Blue nodes represent taxonomic groups only observed in GAZEL-ADN and grey nodes represent taxonomic groups not observed in GAZEL-ADN but present in the HMP data. (b) Proportions of normalized reads mapped to reconstruct 16S rRNA genes showing the 11 different bacterial phyla found for the 39 individuals of GAZEL-ADN.

## Discussion

In this study, we have compared WGS variant calls derived from 39 paired saliva and blood samples. We have demonstrated that a majority of the discordances that are observed between blood and saliva occur in regions of the genome where sequencing is known to be least accurate. We could observe such discordances congregating in these regions due to the relatively large sample size that we had access to for this study. Furthermore, it was clear that recommended quality control measures successfully indicated and excluded a very high proportion of the genotypes that displayed disagreement between blood and saliva.

To fully establish whether the remaining differences after quality control between saliva and blood were remarkable or not, we used a novel method of splitting FASTQ files and creating sequencing pseudo-replicates. This enabled us to confidently establish that the differences between blood and saliva were entirely similar to the differences that one should expect if either an individual’s blood or saliva had been sequenced twice. Thus, while high concordance between sequencing data in blood and saliva has previously been reported [18–20,28,38], we have been able to better contextualise such a result. This has afforded the conclusions that the WGS data generated from the saliva collection kits is of high quality; and that it will be possible to analyse the prospective POPGEN dataset alongside existing datasets (for analyses such as common or rare variant association studies) without fear of harmful batch effects. Technical bias due to differences is technology being used within a study is a non-negligible problem. The large scale of many recent genetic studies and the time required to assemble genomic datasets may result in different sequencing techniques being used at the start and at the end due to evolving guidelines and technology. The herein proposed novel method of pseudo-replication represents a valuable tool for assessing the potential impact of technological bias within a study as it can provide good approximations for benchmarks of wholegenome sequencing reproducibility.

In our study, we have concentrated on appraising the suitability of saliva for WGS in humans for the calling of SNPs and INDELs. In both Yao et al. [18] and Trost et al. [28] the question of the accuracy of the calling copy-number variations (CNVs) from saliva is also discussed where greater discordance (than for SNPs or INDELs) between blood and saliva has been presented. Prospectively, the 39 paired datasets of GAZEL-ADN will provide valuable opportunities for evaluating of the calling of CNVs from saliva and also for similar investigations for structural variants, transposable elements, and mitochondrial variation.

Here, we have also demonstrated that the sets of non-human reads found in saliva carry sufficient information for taxonomical descriptions of the salivary microbiome even when samples were treated for human genome sequencing. We were able to show that the principal groups of bacterial taxa found in our study matched those found in a large public microbial reference dataset. We could also highlight differences between individuals in terms of presence or absence and relative abundance of certain taxa in their salivary microbiome populations. These findings merit further investigation and, as a prospective of this study, we plan to continue to determine a best strategy for extracting salivary microbiome information from WGS data from saliva and to further develop a bioinformatics pipeline for the generation of microbiome data from read data initially generated for human whole-genome sequencing.

This pilot study enables us to have confidence for the use of home-use saliva kits for the proposed new French genetic reference panel POPGEN of the French medical genomics initiative [39,40]. The WGS data generated from saliva will be an equally good approximation of the true genomes of the participants had the project been based on blood collection. As using saliva will also come with considerable benefits in terms of the participation rates and logistics of such a study, it therefore represents a highly attractive data-collection method. We also envisage that it will be possible to study patterns of microbial populations across France in parallel to studies of human genetics via the collection of saliva for sequencing in the POPGEN project.

## Materials and methods

### Data Collection

Sixty individuals from the GAZEL cohort [41] from whom WGS data was already available were selected at random and asked to donate saliva for sequencing. WGS data from saliva of 39 of these individuals were investigated in this study. We refer to this pilot study of 39 individuals as GAZEL-ADN. This study was approved by the *Ile de France XI Ethics Comittee* (approval number 18021) on 8th March 2018 and all the 39 individuals signed an informed consent. The remaining 21 individuals of the original 60 were discarded for the following reasons: 13 individuals did not return their kits, 4 individuals returned their kits too late (outside of a pre-specified time limit of the pilot study) and 4 were removed after failing to meet quality control thresholds for their collected DNA samples. A flow chart of the selection of individuals for the GAZEL-ADN pilot study is included as Supplementary Figure 5. DNA extraction for saliva samples was performed at the CHU Brest, France on a Maxwell (PROMEGA) instrument and using magnetic bead technology. DNA extraction for blood samples was performed at the CEPH in Paris, France on an Autopure (Qiagen) automated system using a salting out method. All libraries for whole-genome sequencing were subsequently prepared using the Illumina TruSeq PCR-free protocol, with an input of 1μg of DNA at the CNRGH in Evry. However, library preparation and sequencing for saliva was performed over two years later than that of blood. The saliva samples were processed using an Illumina NovaSeq 6000 sequencer, while blood samples had previously been processed using an Illumina HiSeqX5 sequencer. These two sequencers have a common underlying technology, and have recently been shown to perform similarly [42]; the main difference being the increased speed and throughput of NovaSeq. Generation of sequencing data was performed using the same bioinformatics pipeline for blood and saliva. Alignment was performed using the software bwa (v.0.7.15) [43] to the human reference genome hs37d5; a human reference genome from genomic build 37 and including dummy contigs for mapping well-known non-human contaminates. This alignment was followed by genotype calling using the GATK Haplotype Caller (v.3.8) [44].

### Comparing blood and saliva WGS call sets

To compare the genotype calls between saliva and blood, the F-measure was used as in [45,46]. The F-measure is the harmonic mean of two other statistics: precision and recall. Precision is equal to TP/(TP+FP) where FP is the number of false positive calls (alternative alleles observed in saliva when only reference alleles were observed in blood) and TP is the number of corresponding true positives (alternative alleles observed in both saliva and blood). Recall is equal to TP/(TP+FN) where FN is the number of false negatives (reference alleles observed in saliva when alternative alleles were observed in blood). If conversely calls from saliva data are to be considered as ‘true’, this interpretation leads to the precision becoming equal to the recall and vice-versa. Hence, the F-measure (which can be calculated per individual or per variant) would not change and so gives an indication of similarity between two sets of genotypes without prioritising either call set. Throughout, we often contrast comparisons of variant call sets calculated before and after applying quality control thresholds. Quality control was performed using VCFProcessor (v.1.0.1) [47] and using an in-house set of thresholds (option QC1078) based on GATK recommendations. Full details of the quality control thresholds applied are given in Supplementary Materials.

### Pseudo-replication

Trost et al., [28] gave a baseline for levels of concordance between blood and saliva from a single external repeated sequencing study for comparisons of sequencing from different biological sources (blood, saliva, and buccal cells). We developed this idea and introduced a novel technique of pseudo-replication. We went back to the original FASTQ files for each individual and split these unordered lists of raw read data in two (retaining reads with their paired ends). This was simply achieved by selecting odd and even groups of four from the line numbers of the two FASTQ files for each individual (one file containing the front end-reads, one file containing the corresponding back-end reads). Naturally, this reduced the depth by 50% to approximately 20× in our study; this lower depth will lead to lower quality in our data, but we should still expect to be able to have well called genotypes at this depth [48]. Hence, each sequencing run provided two pseudo-replicates that are an approximation of results from a replication study at half the depth for each individual. To create our pseudo-replicates, we first applied the quality control program BBTools (v.36.02) [49] to our raw read data. Alignment was then performed using bwa (v.0.7.15) [43] on to the most recent human reference genome GRCh38. Using elprep (v.4.0.0) [50], reads were sorted and both duplicated reads and unmapped reads were removed. Finally, we used the GATK HaplotypeCaller (v.3.8) to call variants with default parameters.

### Exploring the Salivary Microbiome

Illumina shotgun reads were filtered using BBDuk (qtrim=rl, trimq=20, maq=20, minlen=100; BBTools (v.36.02) [49]) to remove Illumina adapters, known Illumina artifacts, phiX, and to quality-trim both ends to Q20. Resulting reads containing more than one ‘N’, or with quality scores (before trimming) averaging less than 20 over the read, or length under 100 bp after trimming, were discarded. Remaining reads were mapped to human indexes with kraken2 (v.2.0.7) [51], discarding all human reads. Small subunit (SSU) rRNA gene sequences were reconstructed from these non-human reads and classified against the SILVA 138 SSU rRNA gene database [52] using phyloFlash (v.3.0) [53]. Fulllength (>70% of the target length) 16s RNA genes were then assembled with SPAdes (v3.13.1) [54] or reconstructed with EMIRGE (v.1.3) [55]. Reads from each of the two subsequently generated libraries were mapped onto the reconstructed sequences with a minimum identity of global alignment of 99% using BBmap (v.36.02) [49] to estimate the relative abundance of each sequence in the respective dataset. Sequences with a coverage statistic of less than 1 were regarded as not present in a sample and removed from further analysis. Taxa abundances were summed at the phylum level and the genus level for specific genera. Figures were realised using the R-packages ‘phyloseq’ [56], ‘taxa’ [57], and ‘metacoder’ [58]. Publically available salivary microbiomes and relevant metadata were downloaded from the HMP (https://www.hmpdacc.org/) in order to provide a comparison dataset and to check quality. Operational taxonomic unit (OTU) representative sequences, OTU tables, and mapping files for 16S rRNA V3-V5 sequencing were downloaded from publically available HMP QIIME Community Profiling datasets (http://downloads.hmpdacc.org/data/HMQCP/otu_table_psn_v35.txt.gz).

## Acknowledgments

We would like to thank all of the members of the GAZEL cohort for their participation in this study. Financial support was obtained from the French Ministry of Research and Innovation for the POPGEN project from the French Medical Genomics Plan. Administrative and regulatory support was provided by Inserm. The extraction and WGS sequencing of blood samples was supported by the Laboratory of Excellence GENMED (Medical Genomics) grant no. ANR-10-LABX-0013 managed by the National Research Agency (ANR) part of the Investment for the Future program. The whole genome sequencing of saliva samples was supported by the CNRGH, CEA. We also thank the CEPH-Biobank team for their involvement in DNA extractions for the GAZEL cohort and the CNRGH teams for their involvement in DNA whole-genome sequencing.

## Data Availability

The complete WGS data generated in this study will be submitted to the French Centralized Data Center of the France Medicine Genomic Plan that is under construction. A synthetic dataset has been generated that allows the replication of our principal results but without a full disclosure of individual level sequencing data. This dataset is currently available on our website http://lysine.univ-brest.fr/~aherzig/BloodSaliva/.

## Author Contributions

AFH carried out analyses pertaining to the comparison of data quality between blood and saliva, and wrote the manuscript. LVS carried out the analyses of salivary microbiomes. EG planned the study and wrote the protocol with CD. All analyses in the study were discussed and decided upon by AFH, LVS, and EG. GL-F acted as project manager regarding the logistics of individual recruitment for this pilot study in conjunction with EG, CD, MG, FL, and MZ. HB managed the blood DNA extractions, LL-R managed the saliva DNA extractions, and AB, RO and JFD contributed to WGS data production. All authors participated in the final redaction of the manuscript.

## Declarations

The authors have no conflicts of interest to declare.

## References

1. the Haplotype Reference Consortium, McCarthy S, Das S, Kretzschmar W, Delaneau O, Wood AR, et al. A reference panel of 64,976 haplotypes for genotype imputation. Nature Genetics. 2016;48: 1279–1283. doi:10.1038/ng.3643

2. Escalona M, Rocha S, Posada D. A comparison of tools for the simulation of genomic next-generation sequencing data. Nat Rev Genet. 2016;17: 459–469. doi:10.1038/nrg.2016.57

3. Zook JM, Chapman B, Wang J, Mittelman D, Hofmann O, Hide W, et al. Integrating human sequence data sets provides a resource of benchmark SNP and indel genotype calls. Nature biotechnology. 2014;32: 246–251. doi:10.1038/nbt.2835

4. Eberle MA, Fritzilas E, Krusche P, Källberg M, Moore BL, Bekritsky MA, et al. A reference data set of 5.4 million phased human variants validated by genetic inheritance from sequencing a three-generation 17-member pedigree. Genome Res. 2017;27: 157–164. doi:10.1101/gr.210500.116

5. Li H, Bloom JM, Farjoun Y, Fleharty M, Gauthier L, Neale B, et al. A synthetic-diploid benchmark for accurate variant-calling evaluation. Nature Methods. 2018;15: 595–597. doi:10.1038/s41592-018-0054-7

6. Hansen T v O, Simonsen MK, Nielsen FC, Hundrup YA. Collection of Blood, Saliva, and Buccal Cell Samples in a Pilot Study on the Danish Nurse Cohort: Comparison of the Response Rate and Quality of Genomic DNA. Cancer Epidemiol Biomarkers Prev. 2007;16: 2072–2076. doi:10.1158/1055-9965.EPI-07-0611

7. Sun F, Reichenberger EJ. Saliva as a source of genomic DNA for genetic studies: review of current methods and applications. Oral health and dental management. 2014;13: 217–222.

8. Brieger K, Zajac GJM, Pandit A, Foerster JR, Li KW, Annis AC, et al. Genes for Good: Engaging the Public in Genetics Research via Social Media. The American Journal of Human Genetics. 2019;105: 65–77. doi:https://doi.org/10.1016/j.ajhg.2019.05.006

9. Haworth S, Mitchell R, Corbin L, Wade KH, Dudding T, Budu-Aggrey A, et al. Apparent latent structure within the UK Biobank sample has implications for epidemiological analysis. Nature Communications. 2019;10: 333. doi:10.1038/s41467-018-08219-1

10. Munafò MR, Tilling K, Taylor AE, Evans DM, Davey Smith G. Collider scope: when selection bias can substantially influence observed associations. Int J Epidemiol. 2018;47: 226–235. doi:10.1093/ije/dyx206

11. Boyd A, Golding J, Macleod J, Lawlor DA, Fraser A, Henderson J, et al. Cohort Profile: the ’children of the 90s’--the index offspring of the Avon Longitudinal Study of Parents and Children. Int J Epidemiol. 2013;42: 111–127. doi:10.1093/ije/dys064

12. Sudlow C, Gallacher J, Allen N, Beral V, Burton P, Danesh J, et al. UK Biobank: An Open Access Resource for Identifying the Causes of a Wide Range of Complex Diseases of Middle and Old Age. PLOS Medicine. 2015;12: e1001779. doi:10.1371/journal.pmed.1001779

13. Quinque D, Kittler R, Kayser M, Stoneking M, Nasidze I. Evaluation of saliva as a source of human DNA for population and association studies. Anal Biochem. 2006;353: 272–277. doi:10.1016/j.ab.2006.03.021

14. Rogers NL, Cole SA, Lan H-C, Crossa A, Demerath EW. New saliva DNA collection method compared to buccal cell collection techniques for epidemiological studies. Am J Hum Biol. 2007;19: 319–326. doi:10.1002/ajhb.20586

15. Thiede C, Prange-Krex G, Freiberg-Richter J, Bornhauser M, Ehninger G. Buccal swabs but not mouthwash samples can be used to obtain pretransplant DNA fingerprints from recipients of allogeneic bone marrow transplants. Bone Marrow Transplantation. 2000;25: 575–577. doi:10.1038/sj.bmt.1702170

16. Hu Y, Ehli EA, Nelson K, Bohlen K, Lynch C, Huizenga P, et al. Genotyping performance between saliva and blood-derived genomic DNAS on the DMET array: A comparison. PLoS ONE. 2012;7: e33968. doi:10.1371/journal.pone.0033968

17. Herráez DL, Stoneking M. High fractions of exogenous DNA in human buccal samples reduce the quality of large-scale genotyping. Anal Biochem. 2008;383: 329–331. doi:10.1016/j.ab.2008.08.015

18. Yao RA, Akinrinade O, Chaix M, Mital S. Quality of whole genome sequencing from blood versus saliva derived DNA in cardiac patients. BMC Medical Genomics. 2020;13: 11. doi:10.1186/s12920-020-0664-7

19. Zhu Q, Hu Q, Shepherd L, Wang J, Wei L, Morrison CD, et al. The impact of DNA input amount and DNA source on the performance of whole-exome sequencing in cancer epidemiology. Cancer Epidemiology Biomarkers and Prevention. 2015;24: 1207–1213. doi:10.1158/1055-9965.EPI-15-0205

20. Kidd JM, Sharpton TJ, Bobo D, Norman PJ, Martin AR, Carpenter ML, et al. Exome capture from saliva produces high quality genomic and metagenomic data. BMC Genomics. 2014;15: 262. doi:10.1186/1471-2164-15-262

21. Abraham JE, Maranian MJ, Spiteri I, Russell R, Ingle S, Luccarini C, et al. Saliva samples are a viable alternative to blood samples as a source of DNA for high throughput genotyping. BMC Med Genomics. 2012;5: 19. doi:10.1186/1755-8794-5-19

22. Bahlo M, Stankovich J, Danoy P, Hickey PF, Taylor BV, Browning SR, et al. Saliva-derived DNA performs well in large-scale, high-density single-nucleotide polymorphism microarray studies. Cancer Epidemiol Biomarkers Prev. 2010;19: 794–798. doi:10.1158/1055-9965.EPI-09-0812

23. Fabre A, Thomas E, Baulande S, Sohier E, Hoang L, Soularue P, et al. Is Saliva a Good Alternative to Blood for High Density Genotyping Studies: SNP and CNV Comparisons? Journal of Biotechnology & Biomaterials. 2012;1: 7. doi:10.4172/2155-952x.1000119

24. Feigelson HS, Rodriguez C, Welch R, Hutchinson A, Shao W, Jacobs K, et al. Successful genome-wide scan in paired blood and buccal samples. Cancer Epidemiology Biomarkers and Prevention. 2007;16. doi:10.1158/1055-9965.EPI-06-0859

25. Paynter RA, Skibola DR, Skibola CF, Buffler PA, Wiemels JL, Smith MT. Accuracy of multiplexed illumina platform-based single-nucleotide polymorphism genotyping compared between genomic and whole genome amplified DNA collected from multiple sources. Cancer Epidemiology Biomarkers and Prevention. 2006;15: 2533–2536. doi:10.1158/1055-9965.EPI-06-0219

26. Rylander-Rudqvist T, Håkansson N, Tybring G, Wolk A. Quality and quantity of saliva DNA obtained from the self-administrated oragene method--a pilot study on the cohort of Swedish men. Cancer Epidemiol Biomarkers Prev. 2006;15: 1742–1745. doi:10.1158/1055-9965.EPI-05-0706

27. Bruinsma FJ, Joo JE, Wong EM, Giles GG, Southey MC. The utility of DNA extracted from saliva for genome-wide molecular research platforms. BMC Res Notes. 2018;11: 8. doi:doi:10.1186/s13104-017-3110-y

28. Trost B, Walker S, Haider SA, Sung WWL, Pereira S, Phillips CL, et al. Impact of DNA source on genetic variant detection from human whole-genome sequencing data. J Med Genet. 2019;56: 809–817. doi:10.1136/jmedgenet-2019-106281

29. Zook JM, Catoe D, McDaniel J, Vang L, Spies N, Sidow A, et al. Extensive sequencing of seven human genomes to characterize benchmark reference materials. Scientific Data. 2016;3: 160025. doi:10.1038/sdata.2016.25

30. Zook JM, McDaniel J, Olson ND, Wagner J, Parikh H, Heaton H, et al. An open resource for accurately benchmarking small variant and reference calls. Nature Biotechnology. 2019;37: 561–566. doi:10.1038/s41587-019-0074-6

31. Sørensen T. A method of establishing groups of equal amplitude in plant sociology based on similarity of species content, and its application to analyses of the vegetation on Danish commons. Kongelige Danske Videnskabernes Selskab. 1948;5 (4): 1–34.

32. Dice LR. Measures of the Amount of Ecologic Association Between Species. Ecology. 1945;26: 297–302. doi:10.2307/1932409

33. Hall NE, Mamrot J, Frampton C, Read P, Steele EJ, Bischof RJ, et al. Blood and saliva-derived exomes from healthy Caucasian subjects do not display overt evidence of somatic mosaicism. Mutation Research/Fundamental and Molecular Mechanisms of Mutagenesis. 2020;821: 111705. doi:10.1016/j.mrfmmm.2020.111705

34. Fricker AM, Podlesny D, Fricke WF. What is new and relevant for sequencing-based microbiome research? A mini-review. Journal of Advanced Research. 2019;19: 105–112. doi:10.1016/j.jare.2019.03.006

35. Human Microbiome Project Consortium. Structure, function and diversity of the healthy human microbiome. Nature. 2012;486: 207–214. doi:10.1038/nature11234

36. Gloor GB, Wu JR, Pawlowsky-Glahn V, Egozcue JJ. It’s all relative: analyzing microbiome data as compositions. Annals of Epidemiology. 2016;26: 322–329. doi:10.1016/j.annepidem.2016.03.003

37. Gloor GB, Macklaim JM, Pawlowsky-Glahn V, Egozcue JJ. Microbiome Datasets Are Compositional: And This Is Not Optional. Frontiers in Microbiology. 2017;8: 2224. doi:10.3389/fmicb.2017.02224

38. Wall JD, Tang LF, Zerbe B, Kvale MN, Kwok PY, Schaefer C, et al. Estimating genotype error rates from high-coverage next-generation sequence data. Genome Research. 2014;24: 1734–1739. doi:10.1101/gr.168393.113

39. Lévy Y. Genomic medicine 2025: France in the race for precision medicine. The Lancet. 2016;388: 2872. doi:10.1016/S0140-6736(16)32467-9

40. Lethimonnier F, Levy Y. Genomic medicine France 2025. Annals of Oncology. 2018;29: 783–784. doi:10.1093/annonc/mdy027

41. Goldberg M, Leclerc A, Bonenfant S, Chastang JF, Schmaus A, Kaniewski N, et al. Cohort profile: the GAZEL Cohort Study. Int J Epidemiol. 2007;36: 32–39. doi:10.1093/ije/dyl247

42. Zhou L, Ng HK, Drautz-Moses DI, Schuster SC, Beck S, Kim C, et al. Systematic evaluation of library preparation methods and sequencing platforms for high-throughput whole genome bisulfite sequencing. Sci Rep. 2019;9: 10383–10383. doi:10.1038/s41598-019-46875-5

43. Li H, Durbin R. Fast and accurate short read alignment with Burrows–Wheeler transform. Bioinformatics. 2009;25: 1754–1760. doi:10.1093/bioinformatics/btp324

44. DePristo MA, Banks E, Poplin R, Garimella KV, Maguire JR, Hartl C, et al. A framework for variation discovery and genotyping using next-generation DNA sequencing data. Nat Genet. 2011;43: 491–498. doi:10.1038/ng.806

45. Telenti A, Pierce LCT, Biggs WH, di Iulio J, Wong EHM, Fabani MM, et al. Deep sequencing of 10,000 human genomes. Proc Natl Acad Sci USA. 2016;113: 11901. doi:10.1073/pnas.1613365113

46. Hwang S, Kim E, Lee I, Marcotte EM. Systematic comparison of variant calling pipelines using gold standard personal exome variants. Scientific Reports. 2015;5: 17875. doi:10.1038/srep17875

47. Ludwig TE, Marenne G, Génin E. VCFProcessor. http://lysine.univ-brest.fr/vcfprocessor/index.html. Accessed 08/10/2020.2020.

48. Kishikawa T, Momozawa Y, Ozeki T, Mushiroda T, Inohara H, Kamatani Y, et al. Empirical evaluation of variant calling accuracy using ultra-deep whole-genome sequencing data. Scientific Reports. 2019;9: 1784. doi:10.1038/s41598-018-38346-0

49. Bushnell B. BBMap short read aligner, and other bioinformatic tools. https://sourceforge.net/projects/bbmap/. Accessed 01/19/2020.2015.

50. Herzeel C, Costanza P, Decap D, Fostier J, Verachtert W. elPrep 4: A multithreaded framework for sequence analysis. PLOS ONE. 2019;14: e0209523. doi:10.1371/journal.pone.0209523

51. Wood DE, Lu J, Langmead B. Improved metagenomic analysis with Kraken 2. Genome Biology. 2019;20: 257. doi:10.1186/s13059-019-1891-0

52. Quast C, Pruesse E, Yilmaz P, Gerken J, Schweer T, Yarza P, et al. The SILVA ribosomal RNA gene database project: improved data processing and web-based tools. Nucleic Acids Research. 2012;41: 590–596. doi:10.1093/nar/gks1219

53. Gruber-Vodicka HR, Seah BKB, Pruesse E. phyloFlash – Rapid SSU rRNA profiling and targeted assembly from metagenomes. bioRxiv. 2019; doi:10.1158/1055-9965.EPI-06-0859.

54. Bankevich A, Nurk S, Antipov D, Gurevich AA, Dvorkin M, Kulikov AS, et al. SPAdes: a new genome assembly algorithm and its applications to single-cell sequencing. J Comput Biol. 2012;19: 455–477. doi:10.1089/cmb.2012.0021

55. Miller CS, Baker BJ, Thomas BC, Singer SW, Banfield JF. EMIRGE: reconstruction of full-length ribosomal genes from microbial community short read sequencing data. Genome Biology. 2011;12: R44. doi:10.1186/gb-2011-12-5-r44

56. McMurdie PJ, Holmes S. phyloseq: An R Package for Reproducible Interactive Analysis and Graphics of Microbiome Census Data. PLOS ONE. 2013;8: e61217. doi:10.1371/journal.pone.0061217

57. Foster ZSL, Chamberlain S, Grünwald NJ. Taxa: An R package implementing data standards and methods for taxonomic data. F1000Res. 2018;7: 272–272. doi:10.12688/f1000research.14013.2

58. Foster ZSL, Sharpton TJ, Grünwald NJ. Metacoder: An R package for visualization and manipulation of community taxonomic diversity data. PLOS Computational Biology. 2017;13: e1005404. doi:10.1371/journal.pcbi.1005404

